# Active fluctuations of axoneme oscillations scale with number of dynein motors

**DOI:** 10.1101/2024.06.25.600380

**Authors:** Abhimanyu Sharma, Benjamin M. Friedrich, Veikko F. Geyer

## Abstract

Fluxes of energy generate active forces in living matter, yet also active fluctuations. As canonical example, collections of molecular motors exhibit spontaneous oscillations with frequency jitter caused by non-equilibrium phase fluctuations. We investigate phase fluctuations in reactivated *Chlamydomonas* axonemes, which are accessible to direct manipulation. We quantify the precision of axonemal oscillations after controlled chemical removal of dynein motors, providing an experimental test for the theory prediction that the quality factor of motor oscillations should increase with motor number. Our quantification reveals specialized roles of inner and outer arm dynein motors. This supports a model in which inner dyneins serve as master pace-makers, to which outer arm dyneins become entrained, consistent with recent insight provided by structural biology.

---

Biological oscillators must function precisely despite inherent biological noise. Theory predicts that oscillator precision depends on the number of coupled pacemaker units [1, 2], and is fundamentally bounded by the rate of energy dissipation [3, 4]. Collections of molecular motors can drive oscillatory motion, and thus represent prime examples of biological oscillators [5–7]. Motor oscillations support active sound detection in hair bundles [8, 9], prevail in muscle sarcomeres [10], contribute to positioning of the mitotic spindle [11, 12], and, drive the regular bending waves of motile cilia [13], which pump fluids and propel swimming microorganisms in a liquid [14–16].

Motile cilia are slender appendages of eukaryotic cells with a typical length of 10 *-* 50 *μ*m. Dynein molecular motors are distributed along the entire length of the *axoneme*, the cytoskeletal core of cilia. The architecture of the axoneme is highly conserved, from protists to plants and humans [17] with a stereotypic arrangement of dynein motors with a length density of approximately 1/nm [18]. Motor activity induces sliding of adjacent doublet microtubules of the axoneme, which is converted into bending as sliding is constrained at the basal end [19]. A dynamic instability, in which motor activity is regulated by the mechanical deformation of the axoneme itself, gives rise to regular bending waves at typical frequencies of 10 *-* 50 Hz. The precise mechanism of motor control is still subject to research [13, 20–22], yet controlled perturbations as performed here may contribute to constrain theoretical models.

The chemo-mechanical cycle of an individual dynein motor couples ATP hydrolysis to the generation of piconewton forces [23]. The transitions from one step of this cycle to the next are stochastic, and should thus result in fluctuating forces at the molecular scale [24– 26]. Remarkable, at the mesoscale, fluctuations in axonemal bending waves are observable by light microscopy [2, 20, 22, 27–32]. Measured noise levels surpass the contribution expected from thermal fluctuations by orders-of-magnitudes. Active fluctuations of motile cilia directly affect biological function. Amplitude fluctuations are predicted to enhance the effective diffusivity of ciliated microswimmers [2, 33]. Phase fluctuations reduce synchronization in collections of cilia and cilia carpets [2, 27– 29, 34], which is important for efficient swimming [27, 35] and fluid pumping [36–39].

A complication of analyzing noise in cilia of intact cells is that slow fluctuations on time-scales of hundreds of beat cycles caused by changes in intracellular ion concentrations [31] superimpose onto the fast fluctuations expected from the stochastic stepping of molecular motors, and that the interpretation of manipulation experiments may be more challenging [32]. In particular, the number of molecular motors in a cilium of given length is fixed and cannot be changed, except genetically.

Noise in ciliary oscillations can be quantified in terms of a quality factor as a dimensionless measure of frequency jitter. A simple mathematical model predicted a linear dependence of the quality factor on motor number [2]. So far, this theory has not been put to an experimental test. Gibbons extracted dyneins from sea urchin sperm and observed a frequency reduction proportional to the duration of motor extraction, but did not assess fluctuations [40].

Here, we establish reactivated axonemes with controlled chemical removal of dyneins as a model system to probe non-equilibrium fluctuations of collective motor dynamics as function of motor number.

## RESULTS

### Measurement of the quality factor in reactivated axonemes

Axonemes extracted from *Chlamydomonas* microalgae can be reactivated in ATP buffer [41], providing unique means for measurement and manipulation, see Fig. 1A. We tracked axoneme centerlines in high-speed video recordings with nano-meter spatial precision [22]. We characterized centerline shapes by their tangent angle profiles *ψ*(*s, t*) as function of arc-length s along the axoneme and time *t* [20]. The power-spectral density of *ψ*(*s, t*), averaged over arc-length *s*, displays a principal Fourier peak at the beat frequency *f*_0_, see Fig. 1B. The square-root of the integrated peak area defines an accurate measure for a beat amplitude *A*. The width of the Fourier peak is indicative of the precision of axoneme oscillations, quantified by a quality factor *Q*. Intuitively, *Q* equals the number of beat-cycles over which the phase de-correlates. A second, alternative definition allows to determine *Q* precisely and robustly [2]. Using PCA, we project each shape on a two-dimensional shape space spanned by its two dominant shape modes, which allows to define a noise-averaged limit-cycle parameterized by a phase variable *φ*, and thus to assign a unique phase *φ*(*t*) to each shape, see Fig. 1C. The correlation function of this phase, *C*(*t*) = ⟨ exp *i*[*φ*(*t*_0_ + *t*) − *φ*(*t*_0_)]⟩, can be fitted by an exponential decay |*C*(*t*)| ≈ exp(− π *f*_0_*t*/*Q*), which defines *Q*, see Fig. 1D. For this typical example of a wildtype (*wt*) axoneme, *Q* ≈ 90.

**FIG. 1.**
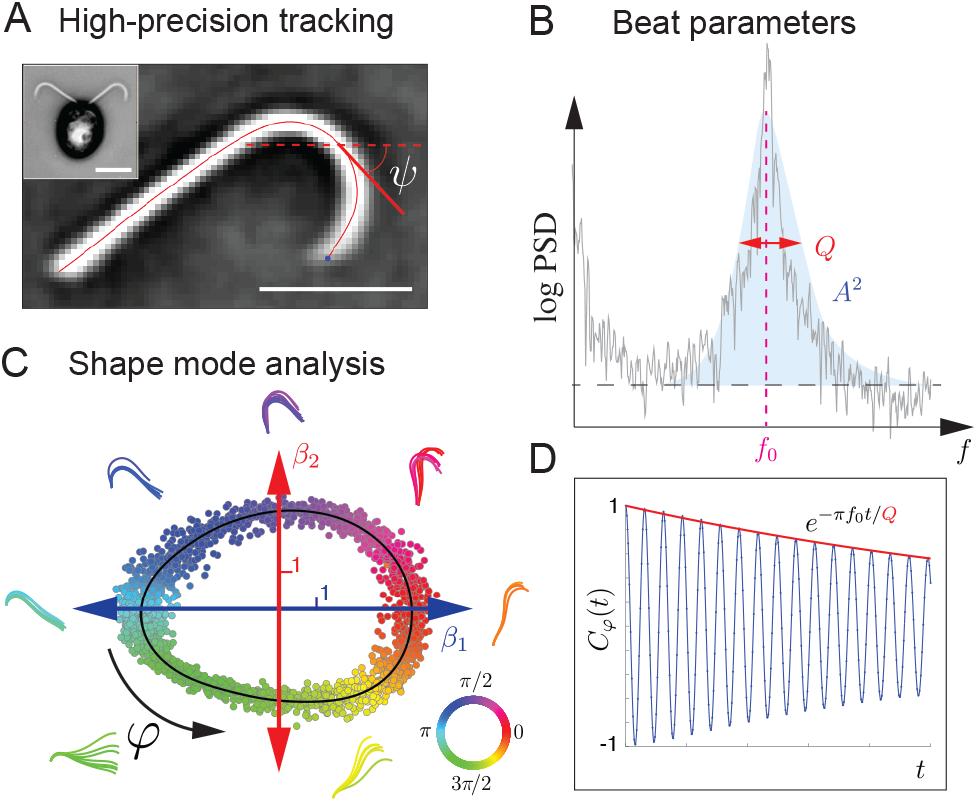
Beating axonemes are noisy oscillators. (*A*) Reactivated axoneme isolated from a *Chlamydomonas* cell (inset). Centerline (red) tracked using FIESTA software [42], characterized by tangent angle *ψ*(*s, t*) as function of arc-length *s* and time *t*. Inverted phase-contrast microscopy. Scale bars are 5 *μ*m. (*B*) The power spectral density of the tangent angle *ψ*(*s, t*) allows to define key parameters of the beat: the position of the fundamental Fourier mode defines the mean beat frequency *f*_0_, while the square root of the integrated power of this peak (highlighted by blue shade) defines a robust measure of beat amplitude *A*. The width of the principal Fourier mode is related to the precision of oscillations, quantified in terms of a quality factor *Q*. (*C*) PCA-projection of waveform data on a shape-space spanned by principal shape modes, where each dot represents a single tracked centerline. To each dot, we assign a beat cycle phase *φ*(*t*), represented by color code, by radial projection onto a limit-cycle of noise-averaged os-cillations (black line). Corresponding axonemal shapes are shown in the respective color. (*D*) The decay of the phase autocorrelation function (red line), which defines the quality factor *Q* of axonemal oscillations

We tested how the quality factor changes for different number of molecular motors using two approaches: first we use a motor mutant (*oda1*, [43]), which is lacking the outer-arm dyneins (OAD) and thus generates motility with only the inner-arm dyneins (IAD). Second, we further reduce the number of dynein motors by biochemical extraction.

### Biochemical extraction of dynein motors

To test the dependence of axoneme oscillations on the number of molecular motors, we developed an assay to biochemically extract dyneins. Axonemes contain two principal types of dynein motors, IAD and OAD, which are repeated in regular arrangement along the entire axonemal length, see Fig. 2A, A’. Salt-extraction with potassium chloride (KCl) is known to extract both IAD and OAD in sea urchin sperm in a concentration-dependent manner [40]. We used SDS-PAGE gels to precisely quantify the amount of remaining IAD and OAD motors (relative to the unextracted *wt* axoneme) in axonemes following extraction, see Fig. 2B. In both *wt* and *oda1* axonemes, the number of motor heads decreases linearly with applied KCl concentration (Fig. 2C and Materials & Methods). For *wt*, we observe a decrease in motor head number of ≈ 20% at the highest KCl concentration of 400 mM. *oda1* axonemes lack all of the OAD motors (corresponding to ≈ 57% of the total *wt* motor heads), and salt-extraction further reduces the number of motor heads to ≈ 30% of the amount present in intact *wt* axonemes. The percentage of axonemes that can be reactivated decreases with the degree of motor extraction (Fig. S3 in SM). Axonemes did not reactivate after salt-extraction at KCl concentrations above 400 mM for *wt*, and 300 mM for *oda1*. Mass Spectroscopy (MS) analysis of the supernatant of salt-extraction (Fig. S2 in SM) revealed that (i) all the major dynein subspecies were extracted, (ii) the composition of different subspecies extracted did not change significantly with KCl salt concentration, (iii) this composition was similar to the stoichiometry in intact axonemes, and (iv) the proportion of all other non-dynein proteins in the salt-extract was less than 0.2 % of total axonemal protein. This validates that the salt-extraction procedure selectively extracts dynein motors. The proportion of IAD and OAD motor heads in the extracted *wt* axonemes, as obtained from the MS data, indicates that the efficiency of extraction of IAD motors was lower in *wt* axonemes as compared to *oda1* axonemes. This could be due to shielding of the IAD motors by the OAD motors when they are present.

**FIG. 2.**
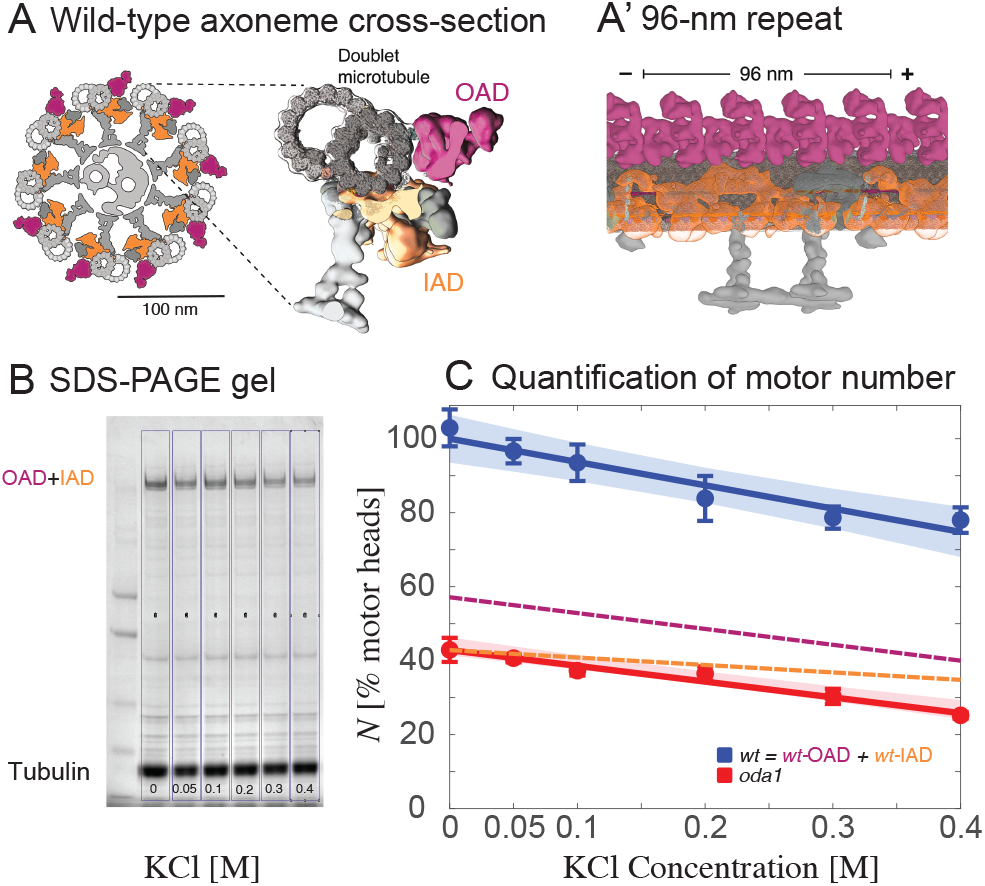
Quantitative extraction of dynein motors. (*A*) Internal structure of the axoneme (modified from [44]): Axonemal cross-section with characteristic 9+2 arrangement of doublet microtubules; the zoom of a doublet microtubule highlights two major motor types: inner arm dyneins (IAD, yellow), and outer arm dyneins (OAD, purple). (*A’*) Stereotypic arrangement of IAD and OAD motors along the length of the axoneme in repeating 96 − nm subunits. The *oda1* mutant lacks all OAD motors. (*B*) Typical example of SDS-PAGE gel of wildtype (*wt*) axonemes treated with KCl with concentrations ranging from 0-0.4 M. Dynein content was quantified relative to tubulin, which is assumed to remain unchanged upon KCl treatment. (*C*) Number of dynein motor heads for both *wt* and *oda1* as function of KCl concentration used in motor extraction (*wt* : blue, *oda1* : red). Dynein-to-tubulin ratios were determined from SDS-PAGE gels and re-scaled using the known stochiometry of dyneins (Fig. S1 in SM). For *wt*, we additionally plot the expected number of OAD (purple) and IAD (orange), with their proportion estimated from mass spectrometry data (Fig. S2 in SM). All values are mean*±* s.e.m. for *n* = 3 independent sets of measurements. Dashed lines represent the 95% confidence intervals.

### Beat frequency is set by both IAD and OAD motor heads

The beat frequency *f*_0_ of reactivated axonemes for both *wt* and *oda1* as function of the total motor head number *N* is shown in Fig. 3A. Beat frequencies without motor extraction are approximately 70 Hz for *wt*, and about 30 Hz for *oda1*, consistent with previous reports [45, 46]. Motor extraction reduces *f*_0_ in a linear fashion similar to previous reports for sea urchin sperm [40]. The collapse of the *wt* and *oda1* data on a single linear relation for beat frequency as a function of motor head number suggests that OAD and IAD equally contribute to setting the beat frequency, irrespective of motor type.

**FIG. 3.**
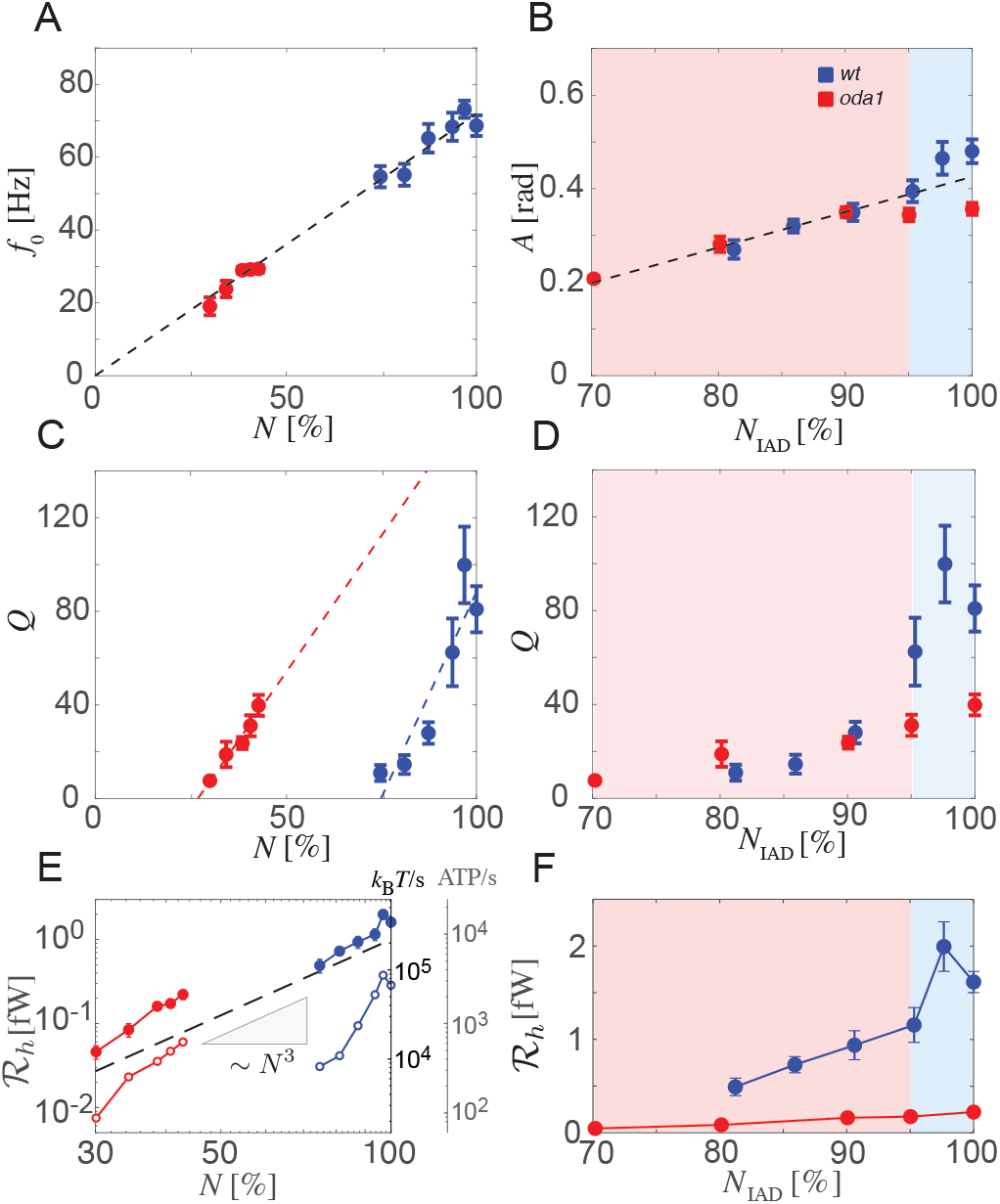
Beating of reactivated axonemes as function of motor number. (*A*) Mean beat frequency *f*_0_ as function of the total number *N* of motor heads (blue: *wt*, red: *oda1*); as well as linear fit of proportionality relation (black dashed). (*B*) Mean beat amplitude *A* as function of the number *N*_IAD_ of IAD motor heads (blue: *wt*, red: *oda1*); as well as linear fit (black dashed). (*C*) Quality factor *Q*, which quantifies the precision of axoneme oscillations, as function of motor head number *N* (blue: *wt*, red: *oda1*); together with separate linear fits (dashed lines). (*D*) Same quality factors *Q*, now as function of the number *N*_IAD_ of IAD motor heads; shaded re-gions *N*_IAD_ *<* 95% (red) and *N*_IAD_ *>* 95% (blue). (*E*) Mean hydrodynamic power output ℛ_*h*_ computed using tracked axonemal beat patterns as function of the total number *N* of motor heads, plotted on a logarithmic scale (closed symbols; blue: *wt*, red: *oda1*). The dashed black line depicts ∼ *N* ^3^. The quality factor measured in the same experiments implies a minimal dissipation rate according to the thermodynamic uncertainty relation [3] (open symbols). Additional vertical axes to the right show alternative energy scales in terms of *k*_*B*_ *T/*s and rate of ATP hydrolysis. (*F*) Same power outputs ℛ_*h*_, now as function of the number *N*_IAD_ of IAD motor heads (error bars smaller than symbols for *oda1*). All values are mean*±*s.e.m.

### Beat amplitude is dominantly set by IAD motor heads

The beat amplitude A of reactivated axonemes for both *wt* and *oda1* as function of the number of IAD motor heads N_IAD_ is shown in Fig. 3B. Remarkably, data for *wt* and *oda1* collapse onto a master-curve, suggesting that IADs predominantly set the beat amplitude. This finding is consistent with previous studies on partial motor mutants [45, 46].

### Quality factor increases with number of motor heads

The quality factor *Q* measures the precision of a noisy oscillator (see Fig. 1C). Fig. 3C shows measured *Q* of reactivated axonemes for both *wt* and *oda1* as function of the total motor head number N. For *wt* axonemes without motor extraction, we find *Q* ≈ 80.9 ± 9.8 (mean ± s.e.m.), consistent with previous estimates for *wt Chlamydomonas* axonemes, based on the width of the principal Fourier peak in the power-spectral density of the tangent angle *ψ*(*s, t*) [46, 47].

For both *wt* and *oda1*, *Q* consistently increases with motor head number N, albeit *oda1* displays lower values with *Q* ≈ 40.7 ± 4.6 (mean ± s.e.m.) without motor extraction.

### OADs affect quality factor only at high IAD densities

Intriguingly, data for *wt* and *oda1* collapse when plotted as function of the number *N*_IAD_ of IAD motor heads, provided *N*_IAD_ ≤ 95% (Fig. 3D). This means that when 5% or more of the IADs are extracted, *Q* is not increased by the additional presence of OADs (red shaded region). In contrast, for *N*_IAD_ > 95%, *Q* increases substantially if OADs are present.

### Impact of ATP concentration on beat parameters

We also studied the beat parameters frequency *f*_0_, beat amplitude *A* and quality factor *Q* as a function of ATP concentration. Consistent with earlier studies, we find that for *wt*, the beat frequency decreases with ATP concentration, following a Michaelis-Menten type relation, see Fig. 4A and [48]. The quality factor displays a similar trend, see Fig. 4C. This ATP-dependency is less pronounced for the oda1 mutant and consistent with a generally higher affinity (lower *K*_*m*_) of IADs to ATP [49]. Thus, OAD activity should be turned down first as the ATP concentration decreases. In contrast, beat amplitude is unaffected by changes in ATP and consistently higher in *wt*, see Fig. 4B. Summarizing our results across treatments, we find that IADs predominantly set beat amplitude and quality factor, while OADs boost beat frequency, see Fig. 4D.

**FIG. 4.**
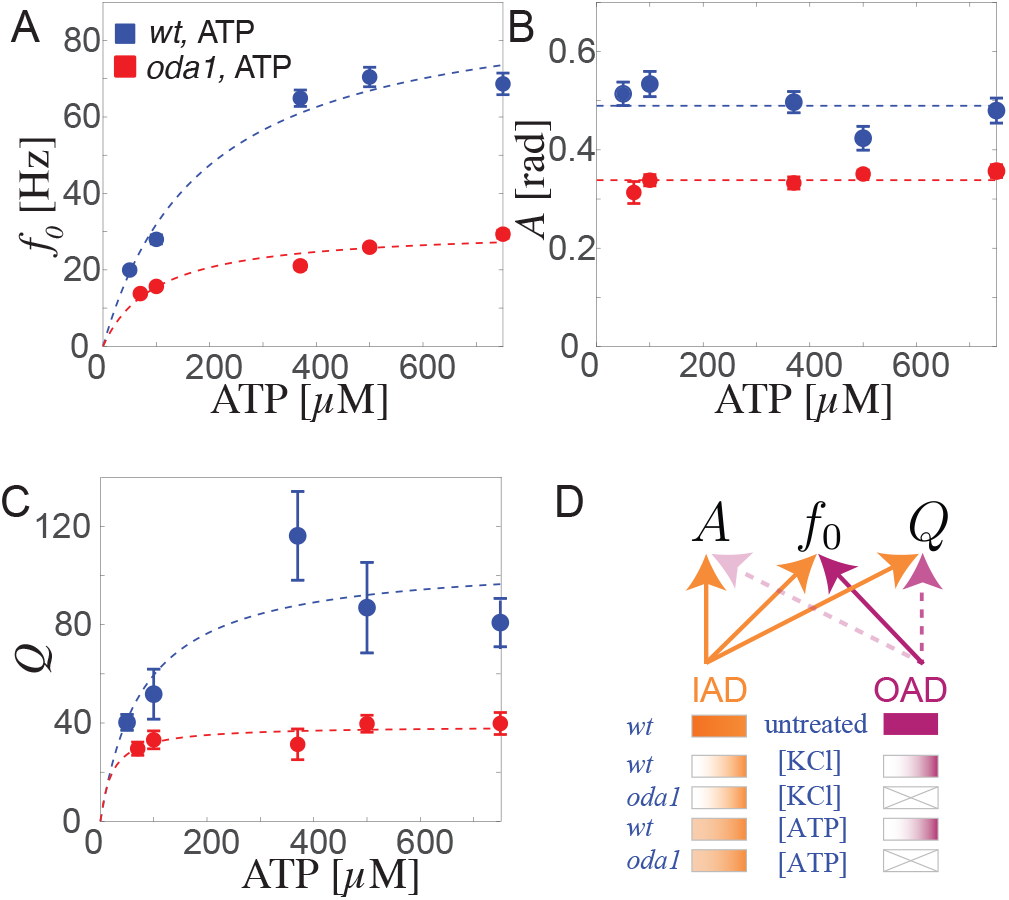
Beating of reactivated axonemes as function of ATP. (*A*) Beat frequency *f*_0_ as function of ATP concentration (blue: *wt*, red: *oda1*). The blue and red dashed curves represent fits of Michaelis-Menten kinetics to the *wt* and *oda1* data, respectively. (*B*) Mean beat amplitude *A* as function of ATP concentration. Dashed lines represent constant fits to the mean values. (*C*) Quality factor *Q* as function of ATP concentration with dashed lines representing Michaelis-Menten fits. All values are mean*±*s.e.m. (*D*) Cartoon summarizing the dependence of the beat parameters *f*_0_, *A* and *Q* on the contribution of IAD and OAD motors manipulated by the treatments considered here (KCl extraction and different ATP concentrations for *wt* and *oda1* axonemes).

### Energetics of axonemal beating

We compute a mean hydrodynamic power output ℛ_*h*_ ≈ 1.6 ± 0.4 fW (mean±s.d., *n* = 15) and ℛ_*h*_ ≈ 0.22 ± 0.16 fW (mean±s.d., *n* = 30) for *wt* and *oda1* axonemes without motor extraction, respectively. This is the time-averaged work exerted by the beating axoneme on the surrounding fluid, which we estimate using resistive force theory [50], see Materials and Methods.

ℛ_*h*_ decreases upon motor extraction and for *oda1* axonemes missing OADs, see Fig. 3E, F (*n* ≥ 10 axonemes for each condition).

The power output ℛ_*h*_ does not scale proportionally to motor number *N* as expected for independent motors, but approximately as ∼ *N* ^3^, (Fig. 3E), consistent with the asymptotic formula ℛ_*h*_ ∼ *A*^2^*f*_0_ [51] and the approximate relationships *A* ∼ *N*_IAD_ and *f*_0_ ∼ *N* reported in Fig. 3A, B, reflecting the cooperative dynamics of molecular motors in the axoneme.

The computed power output is low. For comparison, 1 fW corresponds to the dissipation of approximately 3500 *k*_*B*_*T* during one beat cycle with *f*_0_ = 70 Hz, equivalent to the chemical potential of 150 ATP molecules using the typical energy of 25 *k*_*B*_*T* ≈ 10^*-*19^ J released upon hydrolysis of one ATP molecule [52]. It is likely that a substantial amount of energy released by ATP hydrolysis inside the axoneme is not converted into work exerted on the surrounding fluid, but dissipated internally. Indeed, previous experiments with sperm and *Chlamydomonas*, estimated mechano-chemical energy efficiencies on the order of 10 *-* 40% [53–57] as discussed in [32, 57]. The hydrodynamic power output of *Clamydomonas* cilia beating at 30 Hz was previously estimated as 10 fW [33, 35]. We argue that reactivated axonemes detached from a cell body run idle, and thus their efficiency might be even lower.

### Energetic limits on oscillator precision

Statistical thermodynamics provides an upper bound for the quality factor of active oscillators in terms of their rate of energy dissipation [3], which was recently used to interpret the precision of sperm beating [32]. In our notation, the thermodynamic uncertainty relation (TUR) [3] gives an upper bound on the quality factor *Q* ≤ *Q*_TUR_, where the theoretical limit *Q*_TUR_ = ℛ/(4π *f*_0_ *k*_*B*_*T*) depends on the dissipation rate ℛ and frequency *f*_0_ of the oscillator, and thermal energy *k*_*B*_*T* ≈ 4 zJ at *T* = 20 °C. Fig. 3E re-states this relation as a minimal dissipation rate required by TUR to sustain the measured quality factors of axonemal oscillations.

If we assume an energy efficiency of 20% as previously assumed for intact cells [57], our estimates above correspond to *Q*_TUR_ ≈ 2000 and *Q*_TUR_ ≈ 700 for *wt* and *oda1*, respectively. If there was only one pacemaker per 96-nm repeat (e.g., *IDAf*), and only the energy dissipation of that pacemaker were relevant, these numbers would reduce to *Q*_TUR_ ≈ 100 for both *wt* and *oda1*. This is a variant of the argument of motor coordination put forward by Maggi et al. [32, 58]. On the other hand, it is well possible that the energy efficiency of reactivated axonemes is lower than 20%. If we assume that every motor head hydrolyzes at least one ATP molecule per beat cycle, the energy efficiency would be only 1 %, and hence the estimates for *Q*_TUR_ 20-fold higher.

## DISCUSSION

We measured the precision of a noisy biological oscillator driven by the collective dynamics of molecular motors as function of motor number and ATP concentration, using reactivated *Chlamydomonas* axonemes as model system. We find that the quality factor *Q* of axonemal oscillations increases with the number of inner arm dyneins, consistent with generic theory predictions [2]. Outer arm dyneins increase the quality factor only if 95% of inner arm dyneins are present.

Previous work observed noise in motile cilia and flagella attached to intact cells in terms of a finite-width of the primary Fourier peak in the power-spectral density of ciliary oscillations [20, 46], variations of beat period and noise-induced phase slips in pairs of synchronized cilia [27–29], phase-locking to external oscillatory flow [30], and phase fluctuations in a limit-cycle representation as used here [2, 31, 32]. Our *Q* measurements of isolated wildtype axonemes without motor extraction *Q* ≈ 80 ± 10 are consistent with these previous reports, which estimated *Q* ≈ 25 [28], *Q* ≈ 70 *-* 120 [29], and *Q* ≈ 100 [30] for *Chlamydomonas* cilia. For bull sperm flagella, *Q* ≈ 40 [2] and *Q* ≈ 1 [32] at *T* = 37 °C and *T* ≈ 20 °C, respectively. A reduction of *Q* was observed in shorter cilia [29] and upon oxygen deprivation [32].

Our data indicates specialized roles of inner arm dyneins (IAD) and outer arm dyneins (OAD): IADs serve as the basic driver of the axonemal beat, while OADs are subordinate to IADs to accelerate the beat. This hypothesis is supported by several lines of evidence. First, our study and various earlier reports show that OADs are not essential for ciliary motility [43, 45, 59]. In contrast, motility is strongly affected when specific IADs are removed. Prime examples are *IADf* mutants (*ida1-3*) that have a very irregular beat (*Q* = 15 at 75% N_IAD_), and *pf23* [60], which is missing 76 % of the IADs and, different from other paralyzed mutants, can not be reactivated in low ATP [61]. Second, IADs have been shown to be connected to OADs by specific linkers called outer-inner dynein linkers (OIDLs) [62] and show correlated movement with OADs, suggesting a possible synchronization of the mechano-chemical cycles of both motor types [63]. Taken together, this suggests that OADs might be mechanically controlled by IADs such that their coordinated spatio-temporal activity reduces the variability in local motor activity and increases the quality factor of the beat. Thus, our manipulation experiments suggest that IADs coordinate OAD motors above a critical density.

Our characterization of the axonemal beat for different ATP concentrations is consistent with this hypothesis. Since OAD has a lower affinity to ATP than IAD [49, 64] as discussed in [65], OAD activity should be turned down first upon decrease of ATP concentration, while IAD activity should change less. Indeed, we observe that the beat frequency and the quality factor decrease, while the beat amplitude is unaffected by a decrease in ATP, partially mimicking our results for motor extraction.

Phase noise in a beating axoneme reflects fluctuations in wavespeed of its propagating bending wave. We can abstractly rationalize how the propagation of an activity wave should be affected by motor extraction using a minimal model of coupled excitable motor nodes, see Fig. 5. In short, an active motor node performing a powerstroke is assumed to increase the probability of both transverse and distal neighbor nodes to become activated subsequently, resulting in an activity wave propagating base to tip. The activation of neighboring nodes by an active motor node is stronger if this node comprises OADs and IADs (*wt*) as compared to only IADs (*oda1*). A previously active motor node becomes transiently refractory before it becomes susceptible to activation by neighboring nodes again. Despite strong simplifications, this minimal model qualitatively reproduces the experimental data. We note that alternative models (lacking transverse motor activation, enforced synchronization of IAD and OADs, or a refractory period of motors), could not reproduce the experimentally observed trends. This lets us believe that the model presented here is strongly constrained by the experimental data.

**FIG. 5.**
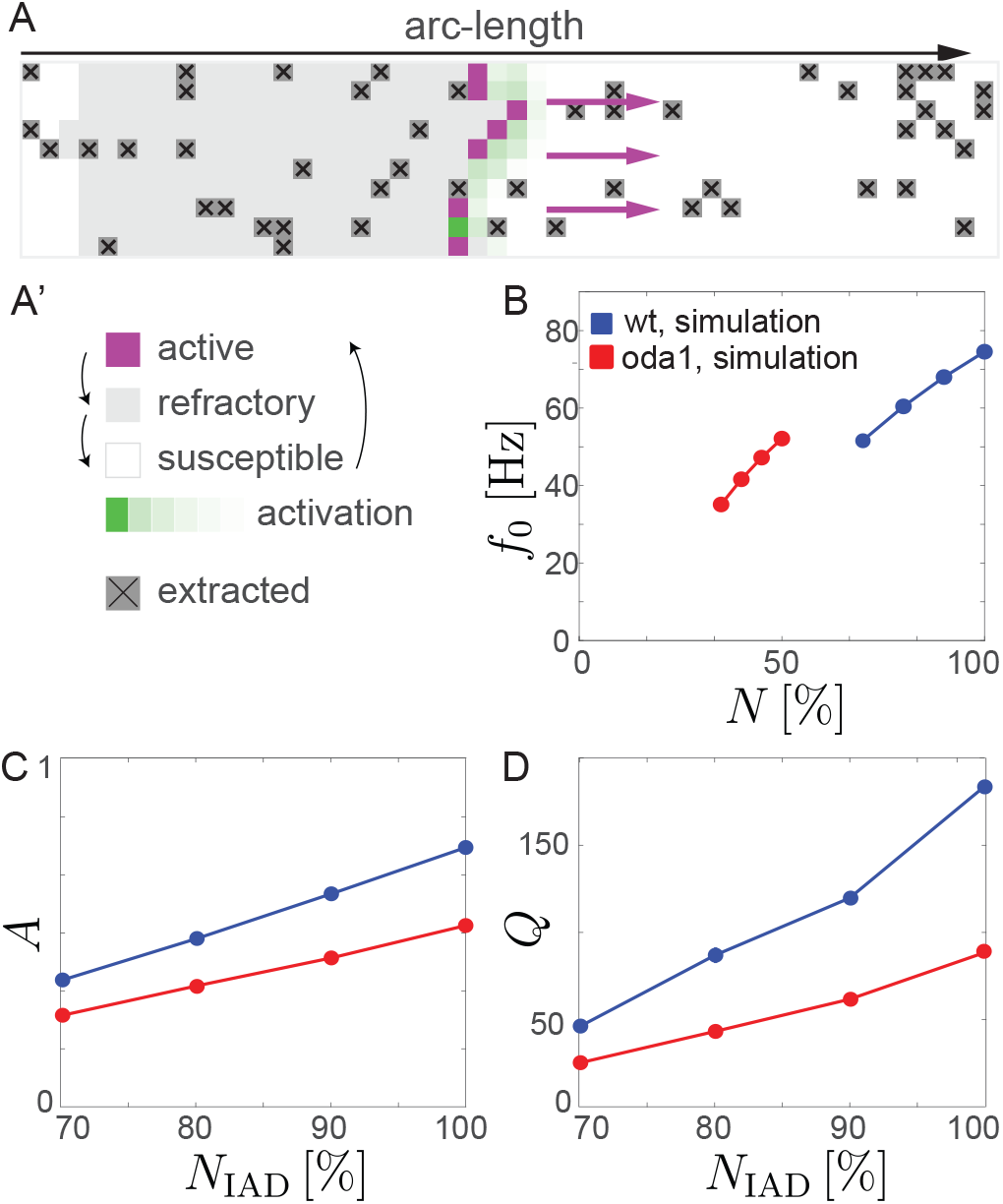
Minimal model of propagating motor activity. (*A,A’*) Activity wave propagating along a square lattice of coupled motor nodes idealizing the axoneme (magenta: active nodes performing power-stroke, light-gray: refractory nodes after power-stroke, white: susceptible nodes, shades of green: activation signal to distal neighbors, dark-gray marked by cross: extracted nodes). (*B*) Emergent frequency *f*_0_ of activity waves (assuming periodic boundary conditions) for partial extraction of motors as function of relative motor number, for lattice consisting of IAD-OAD motor pairs (blue, mimicking *wt*), and lattice of only IAD motor nodes (red, *oda1*). (*C*) Amplitude *A*, defined as time-averaged fraction of active IAD motors as function of total number *N*_IAD_ of IAD motors. (*D*) Quality factor *Q* as function of *N*_IAD_. All values mean of 100 simulation runs, s.e.m. smaller than symbols.

Ciliary motion requires a continuous provision of chemical energy in the form of ATP, which is converted into both work performed on the surrounding fluid and internal heat. We compute a hydrodynamic power output in the femtowatt range for reactivated axonemes, which is lower by an order-of-magnitude than previous estimates for cilia attached to a cell body [32, 33, 66, 67]. Thus, the isolated axonemes investigated here run in idle mode. Correspondingly, the energy efficiency of reactivated axonemes might be lower than that of intact cilia, or, only a fraction of its dynein motors may execute a power-stroke and hydrolyze one ATP molecule per beat cycle. The quality factors of axonemal oscillations measured here are consistent but lower than a theoretic limit imposed by stochastic thermodynamics known as the thermodynamic uncertainty relation [3]. It is possible that the quality factor was not optimized by evolution, with measured values of Q being sufficient for biological function, even if these are much smaller than the theoretic limit.

### Purification and reactivation of axonemes

*Chlamydomonas reinhardtii* strains *wt* (CC-137) and *oda1* (CC-2228) were obtained from *chlamycollection*.*org*. Cells were grown, harvested and axonemes were isolated as in [41]. All reagents were purchased from Merck (sigmaaldrich.com) unless stated otherwise. In brief, cells were grown in a 1 L Tris-Acetate-Phosphate (TAP) medium under constant illumination by LED lightpads (Light Therapy Lamp, 10000 LUX) and air-bubbling at 23 °C, to a final density of 10^6^ cells/ml. Cells were deflagellated using dibucaine, purified through a 25% sucrose cushion, and demembranated in HMDEK buffer (30 mM HEPES-KOH, 5 mM MgSO4, 1 mM dithiothreitol, 1 mM EGTA and 50 mM potassium acetate, at pH 7.4) augmented with 1% (v/v) IGEPAL detergent and 0.2 mM Pefabloc SC protease inhibitor. The demembranated axonemes of 1 l culture were resuspended in 200 *μ*l HMDEK supplemented with 1% (w/v) polyethylene glycol (molecular weight 20 kDa), 28% sucrose and 0.2 mM Pefabloc, and stored at -70 °C.

Axonemes were reactivated in flow chambers (volume 10 *-* 20 *μ*l) built from parafilm strips placed between easy-cleaned microscopy slides and coverslips (22 × 22 mm). The channels of the flow chamber were surface-blocked with casein solution (from bovine milk, 2 mg/mL) for 5 min. Reactivation solution (20 ml) containing HMDEKP buffer (HMDEK plus 1% (w/v) PEG 20.000) with 750 *μ*M ATP and an ATP-regeneration system (1 unit/ml creatine kinase and 5 mM creatine phosphate) was mixed with 1 *μ*l thawed axoneme solution and the channel was sealed with twinsil speed (picodent). For measurements with different ATP concentrations as shown in Fig. 3E, the reactivation buffer contained different amounts of ATP (50, 100, 370, 500, 750 *μ*M for *wt* and 70, 100, 370, 500, 750 *μ*M *oda1*). Axonemes with reduced number of motors were reactivated following a modified protocol (see below). Axoneme reactivation was then observed under the microscope, for a maximum time of 40 min after reactivation.

### Salt-extraction of motors from axonemes

Dynein molecular motors were solubilized and extracted from axonemes using potassium chloride (KCl) salt. The salt extracted axonemes were either reactivated for waveform analysis, or analyzed by SDS-PAGE gels for quantification of motor numbers. The supernatant was analyzed using Mass Spectrometry. Specifically, thawed axonemes (1-2 *μ*l depending on the concentration of axoneme batch) were added to extraction buffer (HMDEK plus 0.2 mM Pefabloc, with KCl added to the desired final concentration) to a total of 50 *μ*l on ice in a 600 *μ*l tube. The KCl concentrations used for extraction were 0 M (unextracted control), 50 mM, 100 mM, 200 mM, 300 mM, and 400 mM. The solution was immediately centrifuged at 10’000 g at 0 °C for 10 min. Extraction was performed on ice and at 0 °C to mitigate the harshness of the treatment. The resulting supernatant containing the extracted motors and excess KCl was removed and either discarded or sent for MS analysis. The resulting pellet consisted of the salt-extracted axonemes. For reactivation, the pellet was directly resuspended in 20 *μ*l of reactivation buffer and flushed into a prepared flow chamber for imaging under the microscope (see below). For motor quantification, the pellet was resuspended in wash buffer (HMDEK plus 0.2 mM Pefabloc) and centrifuged again under the same conditions. This additional centrifugation step was performed to ensure that the samples were sufficiently clean for SDS gel analysis. The resultant supernatant was discarded and the pellet was resuspended in 15 *μ*l of wash buffer. The sample was then prepared for the gel by adding to it 5 *μ*l of 4x SDS sample buffer, followed by heating at 95 °C for 10 min. The entire sample was loaded onto one lane of a 3 - 8% Tris-Acetate gel, which was run at 200 V for 1 hour. The gel was then stained using standard coomassie stain and destained using milliQ water, and then imaged.

### Quantification of motor number in salt-extraced axonemes by SDS-PAGE

Stained and destained SDS-PAGE gels were imaged using a gel-doc system (azure c300) by exposing under visible light using the setting ‘auto exposure time’. In preliminary tests, quantification results were confirmed to be independent of exposure time. Image analysis was performed using the Gel Analyzer tool in the Fiji software suite [68]. Intensity profile plots were generated for gel lanes selected using the rectangular selection tool. The two peaks of interest corresponded to the dynein band at around 500 KDa, and the tubulin band at around 55 KDa. The line selection tool was used to manually draw base lines, which defined the local background. The wand tool was used to measure the area of each enclosed peak (henceforth termed as raw intensity). The raw intensity for both tubulin and dynein was confirmed to scale linearly with the amount of protein loaded onto the gel (Fig. S1 in SM). Assuming tubulin quantity to remain unaffected by the salt-extraction procedure, the ratio of dynein-to-tubulin raw intensity quantified the dynein content normalized to total axoneme content. Mean intensity ratios were calculated from independent sets of gel measurements; for unextracted *wt* and *oda1*, the mean values closely agreed with calculated expectations based on the known stoichiometry of motors in the axoneme (Fig. S1 in SM). A linear regression was fitted to the mean ratios as a function of KCl concentration (weighted by s.e.m.), which was used to compute a look-up table of calibrated intensity ratios.

The percentage of motor heads N relative to the unextracted condition was calculated using these calibrated ratios for both *wt* and *oda1*, respectively. The *oda1* measurements were then rescaled relative to *wt* according to the known stoichiometry of IAD and OAD motor heads (Fig. S1 in SM).

### Analysis of the composition of the salt extract by Mass Spectrometry

Salt concentration of the extracted motor containing supernatant samples was adjusted to < 100 mM for all fractions by dilution with 20 mM NH_4_CO_3_ before insolution digestion. Subsequently samples were digested in solution with trypsin and Lys-C. The digests were desalted with C18 ultra-microcolumns. Elutes were dried in vacuum and stored until analysis at -20°C. The desalted digests were recovered with 3 *μ*l 30% formic acid supplemented with peptide retention time standard at 25 fmol/*μ*l and diluted with 20 *μ*l water. Five microliter were injected for nanoLC-MS/MS analysis with a Dionex 3000 UPLC system hyphenated to a Q-Exactive HF mass spectrometer. The mass spectrometer was operated in DDA mode. The raw data was analysed with Progenesis QIP for feature identification and protein quantification (MI3). Protein identification was performed with the MASCOT. For detailed instrument and software parameters and settings, see (Supplementary Tables S1-5). Common contaminants including human proteins identified in the dataset were deleted. Uncharacterized proteins of mass ∼ 500 kDa were annotated using UniProt and Phytozyme databases, by which all major dynein subspecies were identified [29, 69]. All further analysis was carried out using the raw abundance values of identified proteins. The proportion of IAD to OAD motors was taken to be the mean value of measured proportions in the 200, 300, and 400 mM KCl-extracted samples.

### High-speed imaging of reactivated axonemes

Axonemes were imaged by phase-contrast microscopy set up on an inverted Zeiss Observer Z1 microscope using a 40x Plan-Neofluar NA 1.3 Phase 3 oil lens in combination with a Zeiss oil condenser (NA 1.4) and additional magnification of optovar 2.5x. Illumination was set up using a 100 W tungsten lamp and the Sola SE light-engine (lumencor). Images were acquired using an EoSens 3CL CMOS high-speed camera, with an effective pixel size of 140 nm. Movies consisting of 5000 frames were recorded at the following framerates for different conditions: *wt* KCl series (1000 fps), *oda1* KCl series (690 fps), *wt* ATP series (50 *μ*M: 250 fps, 100 *μ*M: 500 fps, 370 *μ*M:1000 fps, 500 *μ*M: 1500 fps, 750 *μ*M: 1000 fps), *oda1* ATP series (70 *μ*M: 390 fps, 100 *μ*M: 510 fps, 370 *μ*M: 600 fps, 500 *μ*M: 660 fps, 750 *μ*M: 690 fps). Note: the 0 M KCl condition and 750 micromolar ATP condition is the same dataset.

### High-precision tracking of axonemal shapes

Movies were pre-processed using FIJI [68] in order to increase the signal-to-noise ratio. This was done primarily by removing the static background from each frame in the movie. The static background (defined as the image constructed by taking the median intensity value for each pixel over the entire movie) was subtracted from each frame in the movie. Following this, FIJI’s Gaussian blur filter (with setting: Sigma (radius) = 1) was applied for smoothening. This procedure evened out inhomogeneities arising from uneven illumination, remaining dirt particles, etc., increasing the signal-to-background ratio by a factor of 3. The movies were inverted, the contrast was enhanced (using FIJI’s ‘Enhance contrast’ function, with setting: Saturated pixels= 0.35%), and saved as TIFF stacks.

The MATLAB-based filament tracking software FIESTA [42] was used for tracking of the centerline of the axoneme in each movie frame with nanometer precision using a 2D Gaussian fitting algorithm. A manual threshold was set to convert gray-scale movie frames into binary images, and tracking was performed using the following FIESTA settings: full width at half maximum of 800 nm, reducing the fit box size for “tracking curved filaments” by 75%. Using the *x, y* positions of the tracked axoneme centerline, the tangent angle *ψ*(*s, t*) was calculated at 25 equally spaced arc-length points s along the axoneme length for all timepoints *t*. The proximal end *s* = 0 of tracked axonemes was infered from the known direction of wave propagation from the base to the distal tip.

To correct for the global rotation of the swimming axoneme, we subtracted the linear trend of the tangent angle at the center of the axoneme.

### Waveform parameterization by Fourier analysis

To compute the mean beat frequency *f*_0_ and mean beat amplitude *A*, the power-spectral density of the tangent angle *ψ*(*s, t*) was computed as detailed in [22]. The position of the first Fourier peak defines the beat frequency *f*_0_, while the square root of the integrated power of this peak defines the beat amplitude *A*.

### Shape mode analysis

To characterize phase fluctuations, dimensionality reduction using shape mode analysis was used as described [2, 70]. For this, the first tangent angle *ψ*(*s* = 0, *t*) (corresponding to the base) was subtracted from every tangent angle profile. A two-point correlation matrix *M*(*s, s’*) = ∑_*i*_[*ψ*(*s, t*_*i*_) − *ψ*_0_(*s*)][*ψ*(*s*’, *t*_*i*_) − *ψ*_0_(*s’*)] was defined where _0_(s) is the mean shape given by averaging the tangent angle over time. The principal shape modes *ψ*_1_(*s*), *ψ*_2_(*s*) are defined as the eigenvectors corresponding to the two maximal eigenvalues of M, which together accounted for ∼ 98% of the observed variance in the tangent angle data. Each tracked shape was projected onto the shape space spanned by the two principal shape modes as *ψ*(*s, t*_*i*_) ≈ *ψ* _0_(*s*) +, *β*_1_(*t*_*i*_) *ψ*_1_(s) +, *β*_2_(*t*_*i*_) *ψ*_2_(*s*), where the time-dependent coefficients, *β*_1_(*t*),, *β*_2_(*t*) were obtained from a linear least-squares fit. A limit cycle was defined by fitting a closed curve to the point cloud in the shape space formed by the coefficients, *β*_1_,, *β*_2_ assigned to each centerline shape. The limit cycle was parametrized by a phase angle *φ*, defined to increase uniformly along the curve using the renormalization procedure in [71] (using Fourier terms up to the 10^th^ order). A unique phase was assigned to each shape in the point cloud by projecting radially onto the limit cycle. Phase speed fluctuations cause a decay of the phase correlation function, *C*(*t*) = ⟨exp[*i*[*φ*(*t*_0_ + *t*) − *φ*(*t*_0_)]]⟩. The phase diffusion coefficient *D* was determined by fitting an exponential decay to the correlation function according to |*C*(*t*)| ≈ exp(− *D*|*t*|), using values of the correlation function up to a maximal lag time *t*_max_. This maximal lag time *t*_max_ was adapted to the decay time for the different experimental conditions such that *C*(*t*_max_) ≈ 0.5. The quality factor *Q* is then defined as *Q* = 2π *f*_0_/(2*D*).

### Other data filtration

Commonly, only a subset of extracted axonemes can be reactivated; while the rest does not resume beating or beats with abnormal waveforms, likely due to mechanical damage during the extraction procedure.

To exclude axonemes with incomplete reactivation, we employed an automated selection method. First, tracked datasets for which no principal Fourier peak could be detected in the power-spectral density of the tangent angle (using a frequency cutoff of 5 Hz) were discarded (about 13 % of tracked datasets). Second, tracked datasets with abnormal waveforms were excluded (less than 4 % of tracked data sets). For this, we defined an arc-length dependent phase *φ*(s) from the Fourier transform of the tangent angle *ψ*(*s, t*) for each arc-length position s. We then determined the deviation of these phase profiles *φ*(*s*) from mean phase profiles computed for either *wt* or *oda1* axonemes at 0 M KCl, respectively, and discarded tracked data sets for which the root-mean-squared-error (RMSE) of this deviation exceeded 0.3 (Fig. S4 in SM).

Additionally, tracked datasets can contain temporal gaps of missing frames, e.g., if axonemes transiently get out-of-focus. We discarded datasets for which more than 25% of frames were missing (less than 3% of all tracked datasets). Using synthetic data with random deletion of frames, we confirmed that a fraction of 25% of missing frames results in an error of less than 10% in determining the phase diffusion coefficient *D*_0_.

### Computation of hydrodynamic power output

For each tracked axoneme, we computed the hydrodynamic power output ℛ_*h*_ of axonemal beating. We first determined a noise-averaged beat pattern by fitting a Fourier series in *φ* (up to third harmonics) to the tangent angle time-series *ψ*(*s, t*) at each arc-length position *s*. We discarded a small fraction of data sets (3%) for which the mean sum-of-squared-errors of this fit exceeded 1 rad^2^. We then used resistive force-theory [50] to compute the phase-dependent rate of hydrodynamic power Output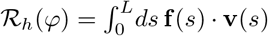, where **f** (*s*) denotes the line density of hydrodynamic friction forces, and **v**(*s*) the velocity of the axoneme at each arc-length position relative to the laboratory frame. We used the friction coefficients *ξ*_‖_ = *ρ* 0.69 10^−3^ pN s/*μ*m^2^ *ξ*_⊥_ = 1.81 *ξ*_‖_ obtained in [72] by a fit to bull sperm data, with a correction factor *ρ* = 1.0/0.7 to account for the different viscosities of water at 36°C [72] and 20°C (this study).

### Minimal motor model

To illustrate the impact of motor extraction on the regularity of emergent cilia dynamics, we devised a conceptual model of a propagating activity wave using a discrete excitable medium consisting of coupled motor nodes. The power-stroke of an IAD motor is assumed to activate motors at the same and the next arc-length position by increasing their powerstroke rate *r*. Each IAD motor is connected to an OAD motor that strikes synchronously, and contributes an equal increase to the neighboring IADs’ rate *r*. After a stroke, each IAD becomes refractory for a transient period of time. This coupling reflects the mechanical coupling of IAD and ODA through outer-inner dynein linkers (OIDLs) [62], which were suggested to be responsible for mechanical coupling and coordination of IAD and OAD [63]. By imposing periodic boundary conditions (akin to a scenario where a bending wave reaching the distal tip of the axoneme initiates a new wave via hydromechanical forces), we obtain a noisy oscillator. Fig. 5 displays frequency *f*_0_, amplitude *A* of IAD powerstrokes, and quality factor *Q* as function of motor fraction. Specifically, the strike rate *r*_*i,j*_ of the IAD motor node at position (*i, j*) obeys

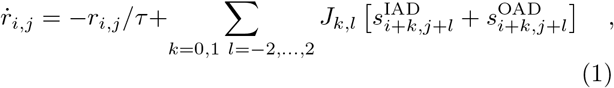

where *τ* is a relaxation time, *J*_*k,l*_ are coupling constants, and 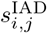 and 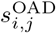 are the powerstroke spike trains of IAD and OAD motor nodes, respectively. Powerstrokes of IAD motor nodes are determined as independent inhomogeneous Poisson processes with rates *r*_*i,j*_. Likewise, extraction of motor nodes is assumed independent. OAD motor nodes strike synchronously with their corresponding IAD motor node, yet become extracted independently. After a power-stroke, an IAD motor node is inactive for a recovery period of duration *t*_refrac_. Parameters *τ* = 2 ms, *J*_1,*±*2_ = 1/(4*τ*), *J*_1,*±*1_ = 1/(2*τ*), *J*_1,0_ = 1*τ, J*_0,*j*_ = 10 *J*_1,*j*_, *J*_*i,j*_ = 0 otherwise, *t*_refrac_ = 4 *τ*, simulation time 10^3^ *τ*.

## Supporting information

Supplementary Materials contain additional details on methods

## Data, Materials and Software Availability

Raw image data and axoneme centerline data are made available upon request. All other data and analyses are included in the main text or in the supplementary materials.

## Acknowledgments

We thank S. Diez for funding and discussions, M. Striegler for discussions, M. Gentzel for Mass Spec experiments. AS was supported by the German Academic Exchange Service (DAAD) and DIGS-BB, TU Dresden. BMF was supported by the Deutsche Forschungsgemeinschaft (DFG, German Research Foundation) under Germany’s Excellence Strategy - EXC-2068-390729961, as well as through a Heisenberg grant (421143374).

## Notes

### Competing Interest Statement

The authors have declared no competing interest.

