## Supplementary Materials contain additional details on methods for "Active fluctuations of axoneme oscillations scale with number of dynein motors"

### S1: Gel analysis additional plots

In order to use the ratio of intensities of dynein to the tubulin band in SDS-PAGE gels as a measure of relative dynein content, it was necessary to confirm that both dynein and tubulin intensities scale linearly with the amount of protein in the concentration range used for experiments. For this we used serial dilutions of *wt* axonemes in the range of 50-200% of the typical concentration used. Figure S1A shows the SDS-PAGE gel for the dilution series. Panels S1B and C show the raw intensity measured using the FIJI gel analyzer tool for dynein and tubulin, respectively. In both cases, the intensity is a well described as a linear function of axoneme concentration.

Figure S1D displays the raw dynein-tubulin ratio from independent gel measurements as a function of KCl concentration used for extraction, with a linear function fitted to the mean values. The measured datasets were selected to be internally consistent, such that $R^2 > 0.7$ for a linear fit to each individual dataset; this was true for $n = 3$ (displayed here) out of 4 gel datasets for both *wt* and *oda1*). The plot also shows estimates for the dynein-to-tubulin ratios calculated purely on the basis of the structural knowledge of the axoneme. Using the well-defined 96 nm repeat of the axoneme, the ratios were estimated as follows:
The axoneme consists of nine microtubule doublets (10+13 protofilaments each) and the central pair (13 protofilaments each), which together comprise a total of 233 protofilaments. Within the 96 nm repeat with a tubulin dimer every 8 nm, this corresponds to 2796 tubulin dimers, or 5592 tubulin monomers in total.

In order to count the total number of dynein motor heads, we consider that there are 4 trimeric (three-headed) OAD motors present on 8 out of the 9 doublets, and 6 monomeric plus one dimeric (dual-headed IAD f) IAD motors present on all 9 doublets. This corresponds to 96 OAD and 72 IAD motor heads, totalling to 168 motor heads for *wt*, and 72 motor heads for *oda1* axonemes per 96 nm repeat. Considering a typical *Chlamydomonas* axoneme of length 10 $\mu$m, this corresponds to a total of 17500 motor heads for *wt*, and 7500 IAD motor heads for *oda1* axonemes. Having counted the absolute numbers of dynein and tubulin units present in the axoneme, we next consider a key assumption. Along with the quantity of protein present, the gel intensity would also be proportional to the size of the protein. This is because larger peptides would contain proportionately greater numbers of the positive amino acid residues to which the Coommassie dye molecules bind. Therefore, taking the molecular masses of the dynein motor head heavy chain and tubulin monomers into account, we have

$$\text{Expected ratio}_{wt} = \frac{168 \text{ (dynein motor heads)} \times 500 \text{ (kDa)}}{5592 \text{ (tubulin monomers)} \times 55 \text{ (kDa)}} \approx 0.27$$

as well as

$$\text{Expected ratio}_{oda1} = \frac{72 \text{ (dynein motor heads)} \times 500 \text{ (kDa)}}{5592 \text{ (tubulin monomers)} \times 55 \text{ (kDa)}} \approx 0.12 \quad .$$

These calculated expectations agree closely with the measured mean dynein-tubulin intensity ratios, as seen in Figure S1D.

In order to go from these intensity ratios to the number of motor heads relative to the intact axoneme, we scaled the calibrated ratios (obtained from the linear fit) for *wt* and *oda1* to the unextracted case for both respectively. Finally, the entire *oda1* data was re-scaled to reflect the fact that the intact *oda1* axoneme, containing only IADs but not OADs, has only 44% of the motor heads present compared to intact *wt*.

### S2: Mass Spec Analysis additional plots

### S3: Quantification of number of beating axonemes

In order to quantify the effect of KCl extraction on the number of reactivating axonemes in the population, we examined large field-of-views of extracted and reactivated axonemes. Under imaging conditions of 40x magnification without additional optovar, covering an area of 496 × 496 pixels with effective pixelsize of 350 nm/pixel, we recorded movies at 1000 fps with total length of 1000 frames. We manually counted and classified the observed axonemes as properly beating, non-beating, or an intermediate state based on the swimming speed, amplitude and waveform as noted visually. Figure S3A reports the count of axonemes of each class as a function of KCl concentration, while Figure

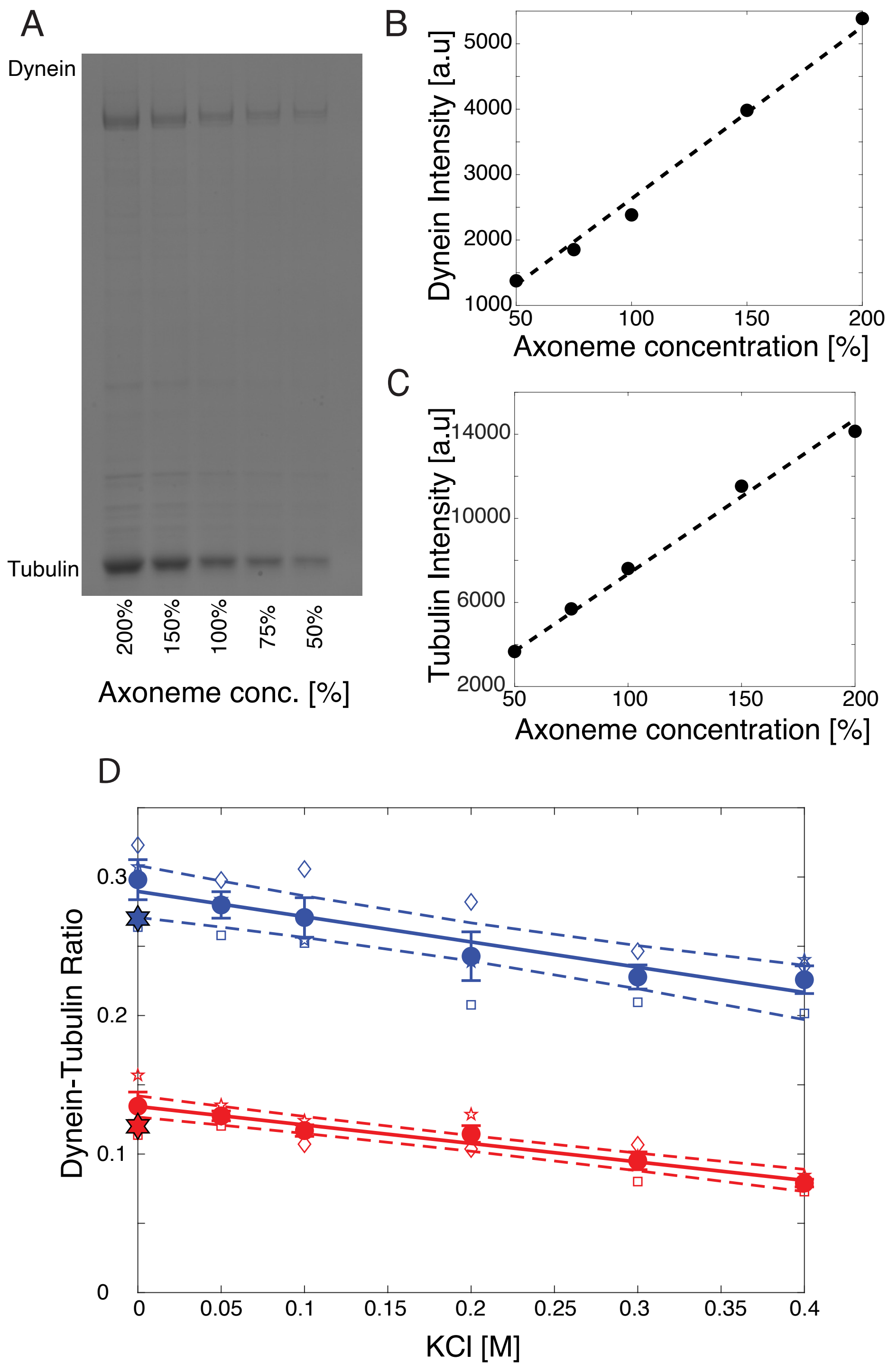


FIG. S1. **SDS-PAGE gel analysis for quantification of motor number**. **A.** SDS-PAGE gel of *wt* axonemes of different relative concentrations. 100% corresponds to the concentration used for all experiments. **B.** Dynein band intensity as function of axoneme concentration, along with linear fit of proportionality relation (black dashed, slope = 26.3). **C.** Tubulin band intensity as function of axoneme concentration, along with linear fit of proportionality relation (black dashed, slope = 73.5). **D.** Dynein-to-tubulin intensity ratio determined from SDS-PAGE gels as function of KCl concentration used in motor extraction (*wt*: blue, *oda1*: red). All values are mean±s.e.m. for $n = 3$ independent sets of measurements (open symbols). Solid lines represent linear fits to the mean intensity ratios (weighted by s.e.m), which correspond to the calibrated intensity ratios used for further analysis. Dashed lines denote 95% confidence intervals. For comparison, we additionally plot the expected dynein-tubulin ratios for *wt* and *oda1* calculated based on the known axoneme structure (blue and red hexagrams).

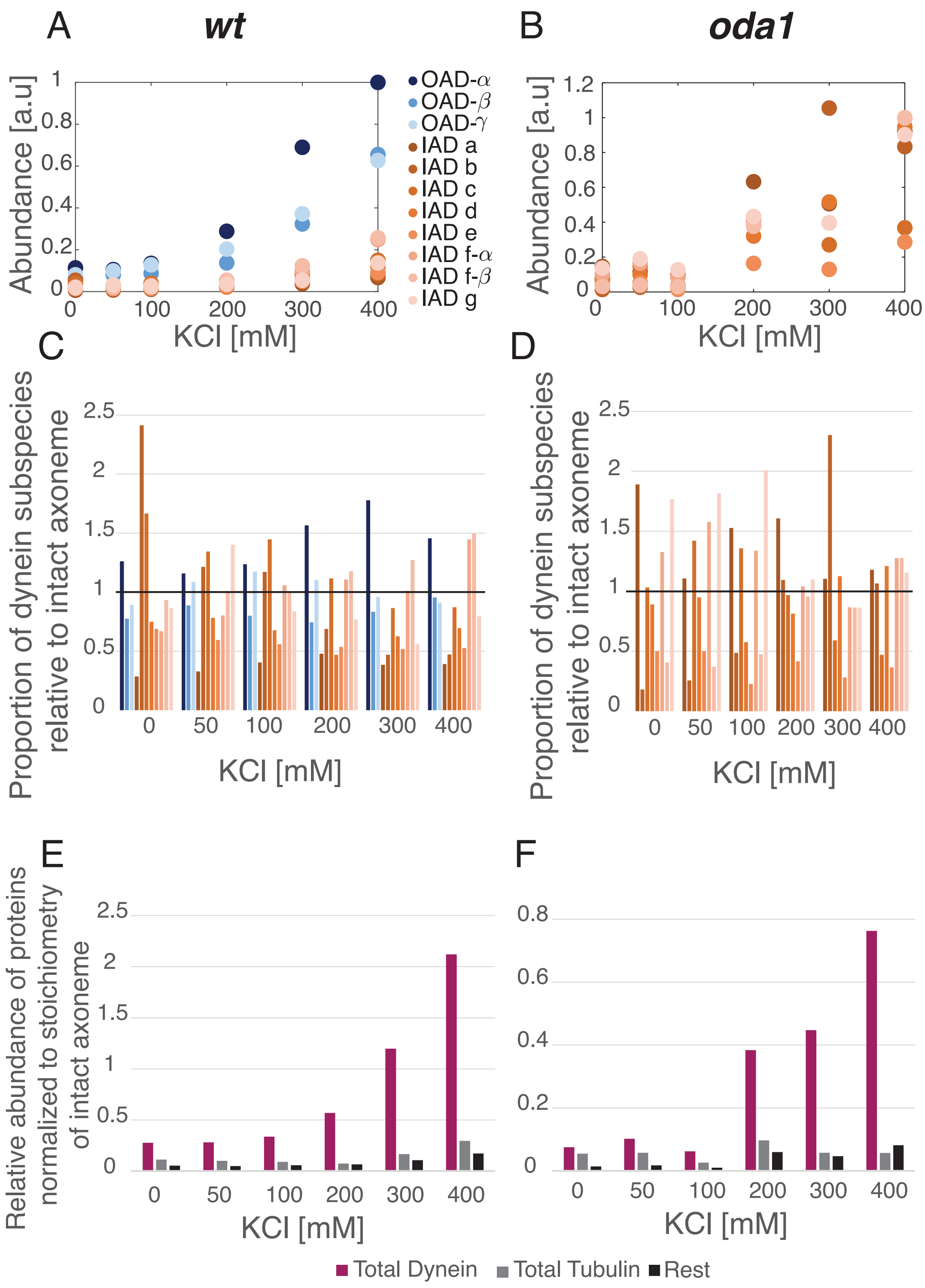


FIG. S2. **Mass spectrometry of supernatant from KCl salt extraction of axonemes. A., B.** Relative abundance of major dynein subspecies in the supernatant of KCl-extracted *wt (A.)* and *oda1 (B.)* axonemes as function of KCl concentration. **C., D.** Proportion of major dynein subspecies in the supernatant of KCl-extracted *wt (C.)* and *oda1 (D.)* axonemes, relative to the stoichiometry of dynein subspecies present in intact *wt* and *oda1* axonemes respectively. Proportion of 1 (black line) indicates that the proportions of extracted motors exactly matches that of intact axonemes. **E., F.** Relative abundance of the total dynein content, total tubulin content, and summed total of all other proteins identified in the supernatant of KCl-extracted *wt (A.)* and *oda1 (B.)* axonemes as function of KCl concentration, normalized by the estimated stoichiometry of proteins present in intact axonemes.

S3B expresses these numbers as relative fractions. It can be seen that the number of beating axonemes decreases drastically at 300 mM, and especially 400 mM KCl. This could be due to the extraction of a critical protein required for beat propagation or the central pair extraction. It is important to note that the data shown in Fig. 3A-D for reactivated axonemes treated with 300 mM and 400 mM KCl comes from sampling from this very small sub-population of beating axonemes.

### A. S4: Data analysis and filtration

Temporal gaps within tracked datasets occur because axonemes can transiently be out-of-focus casing a breakdown of the tracking algorithm in the presence of high imaging noise. This leads to the computed phase $\varphi$ to have improper time-labeling, resulting in incorrect measurements of quality factor $Q$.
Since PCA decomposition depends only on the variation in data independent of time, we compute shape modes from the tangent angle data irrespective of any temporal gaps. In order to keep the time-labeling of the computed phase $\varphi$ consistent, NaN values were filled in corresponding to the position of missing frames. The phase correlation $C(t) = \langle \exp i[\varphi(t_0 + t) - \varphi(t_0)]\rangle$ was then computed with the mean taken by ignoring NaN values.
Using synthetic data with systematic deletion of frames, we tested the effect of missing frames in determining phase diffusion coefficient $D_0$. Synthetic data of defined noise strength was generated as described in [70]. In brief, shape modes were defined as $\psi_1(s) = cos(2\pi s/\lambda)$ and $\psi_2(s) = sin(2\pi s/\lambda)$, and the phase was set as $\dot{\varphi} = \omega_0 + \xi(t)$, where $\xi(t)$ denotes Gaussian white noise with $\langle \xi(t)\xi(t')\rangle = 2D_0\delta(t - t')$. In order to replicate missing frames, a total of 25% of the generated datapoints were deleted under three different test conditions- *1)* uniformly randomly throughout the whole time series, *2)* as a single continuous chunk representing a 'major gap', and *3)* as five chunks of equal size representing 'medium gaps'. Figure S4 compares the $D_0$ as computed using shape mode analysis for the different test conditions. The mean computed $D_0$ remains approximately equal for all test cases. Uniformly random missing frames result in a standard deviation of less than 1.5 %, and the medium or major gap(s) result in a standard deviation of less than 10 % of the value measured in the case of no missing frames. We thereby conclude that datasets with a fraction of upto 25 % missing frames result in an error of less than 10 %, which we deem to be acceptable for analysis.

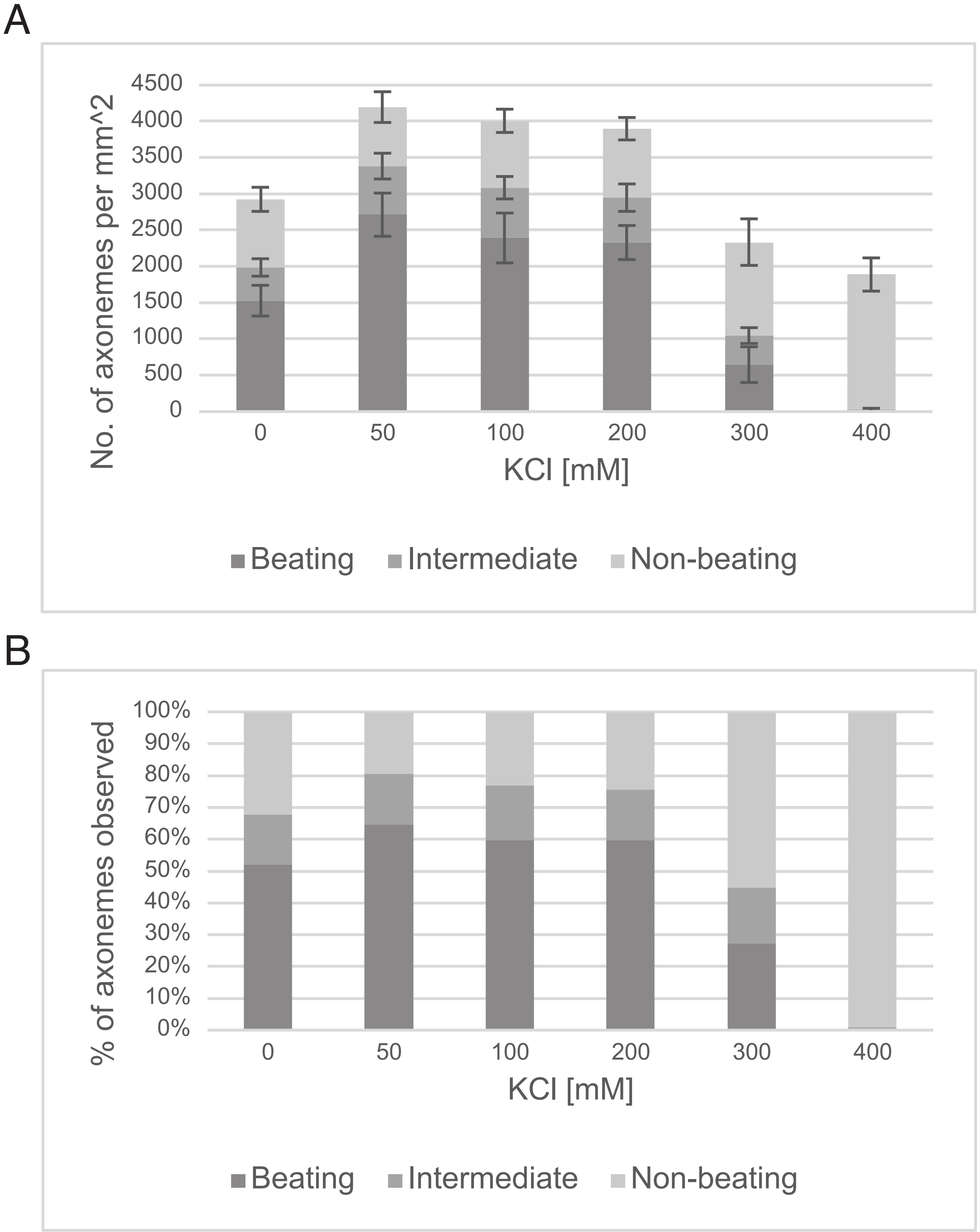


FIG. S3. **Activity of axonemes as function of KCl extraction. A.** Counting of axonemes in a $3.01 \times 10^4\,\mu$m field of view as a function of KCl concentration used for extraction. Observed axonemes were manually classified by visual inspection as beating, non-beating, or intermediate. Counts are reported as mean±s.d. for $n = 10$ different field of views in a single experiment. **B.** Relative fractions of beating, non-beating and intermediate axonemes as a function of KCl concentration determined from the absolute counts in panel A.

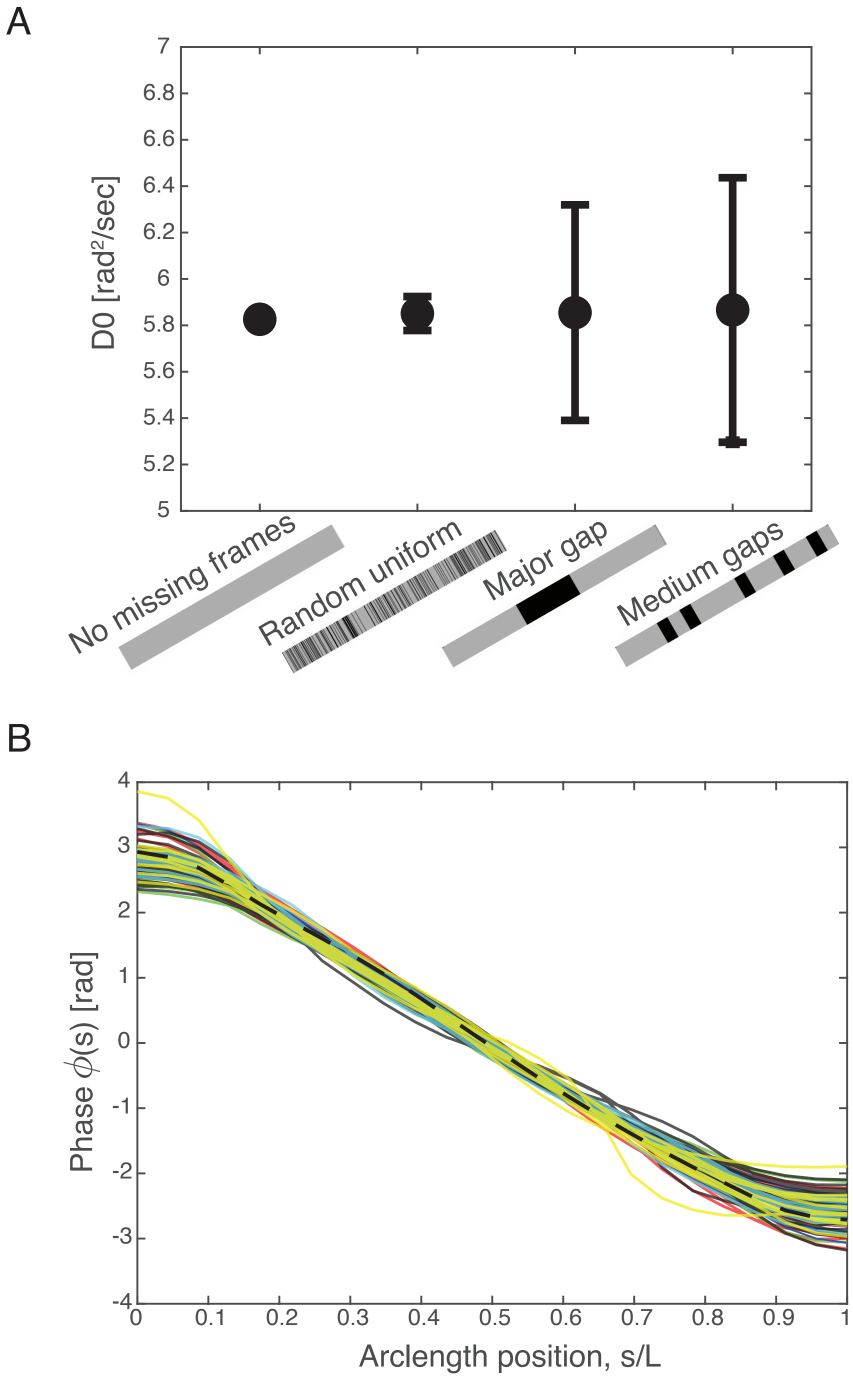


FIG. S4. **Strategies for data processing and selection. A.** Comparison of the phase diffusion coefficient $D_0$ as computed using shape mode analysis for synthetic data under the following conditions- a control case without any missing frames, 'Random uniform' consisting of uniformly random missing frames throughout the time series, 'Major gap' consisting of a single continuous chunk of missing frames positioned randomly along the time series, and 'Medium gaps' consisting of five equal non-overlapping chunks of missing frames positioned randomly along the time series. Illustrations below axis labels represent example distributions of the intact (grey regions) and missing (black regions) frames in each case. Synthetic data was constructed with an original length of 5000 frames, setting $D_0 = 6\text{s}^{-1}$, and a total fraction of 25% of missing frames in the three test cases. Values displayed are mean ± s.d for $n = 25$ randomly generated synthetic datasets.**B.** Using phase-profiles to determine well-behaved waveforms. Phase profiles of *wt* KCl dataset (0M: red, 50 mM: green, 100 mM: blue, 200 mM: black, 300 mM: cyan, 400 mM: yellow) are plotted as function of arclength, along with the mean phase profile for *wt* 0M (black dashed).

| *wt* KCl | Conditions: | 0M | 50 mM | 100 mM | 200 mM | 300 mM | 400 mM | **TOTAL** |
|---|---|---|---|---|---|---|---|---|
| | **Initial** | 16 | 22 | 13 | 26 | 16 | 14 | 107 |
| Data excluded | Fourier-Peak missing | 1 | 0 | 0 | 0 | 0 | 1 | 2 |
| | MSE of C(t) | 0 | 0 | 0 | 0 | 1 | 1 | 2 |
| | Missing frames criterion | 0 | 1 | 0 | 0 | 0 | 0 | 1 |
| | Phase-profile | 0 | 3 | 0 | 2 | 0 | 2 | 7 |
| | **Final** | 15 | 18 | 13 | 24 | 15 | 10 | 95 |

| *oda1* KCl | Conditions: | 0M | 50 mM | 100 mM | 200 mM | 300 mM | | **TOTAL** |
|---|---|---|---|---|---|---|---|---|
| | **Initial** | 34 | 29 | 23 | 14 | 6 | | 106 |
| Data excluded | Fourier-Peak missing | 3 | 0 | 0 | 3 | 3 | | 9 |
| | MSE of C(t) | 0 | 0 | 0 | 0 | 0 | | 0 |
| | Missing frames criterion | 2 | 2 | 3 | 1 | 0 | | 8 |
| | Phase-profile | 0 | 0 | 0 | 1 | 0 | | 1 |
| | **Final** | 29 | 27 | 20 | 9 | 3 | | 88 |

| *wt* ATP | Conditions: | 50 $\mu$M | 100 $\mu$M | 370 $\mu$M | 500 $\mu$M | 750 $\mu$M | | **TOTAL** |
|---|---|---|---|---|---|---|---|---|
| | **Initial** | 11 | 11 | 11 | 11 | 16 | | 60 |
| Data excluded | Fourier-Peak missing | 0 | 0 | 0 | 0 | 1 | | 1 |
| | MSE of C(t) | 0 | 0 | 0 | 0 | 0 | | 0 |
| | Missing frames criterion | 0 | 0 | 0 | 0 | 0 | | 0 |
| | Phase-profile | 0 | 0 | 0 | 0 | 0 | | 0 |
| | **Final** | 11 | 11 | 11 | 11 | 15 | | 59 |

| *oda1* ATP | Conditions: | 70 $\mu$M | 100 $\mu$M | 370 $\mu$M | 500 $\mu$M | 750 $\mu$M | | **TOTAL** |
|---|---|---|---|---|---|---|---|---|
| | **Initial** | 24 | 34 | 27 | 66 | 34 | | 185 |
| Data excluded | Fourier-Peak missing | 8 | 14 | 6 | 15 | 3 | | 46 |
| | MSE of C(t) | 0 | 0 | 0 | 0 | 0 | | 0 |
| | Missing frames criterion | 1 | 1 | 1 | 0 | 2 | | 5 |
| | Phase-profile | 6 | 1 | 0 | 1 | 0 | | 8 |
| | **Final** | 9 | 18 | 20 | 50 | 29 | | 126 |

FIG. S5. Number of datasets analyzed and excluded for each condition.

| Instrument / Parameter | Value | Comments |
|---|---|---|
| **Q-Exactive HF** | ThermoScientific, Bremen, Germany | DDA-Mode (positive ion mode) |
| Tune 2.9 | ThermoScientific, Bremen, Germany | MS operating software |
| Xcalibur / Foundation 3.0 | ThermoScientific, Bremen, Germany | Data visualization & interpretation |
| **MS1** | | |
| Polarity | Positive | |
| Resolution | R120000 at m/z 200 | |
| AGC | 3x 10E6 | |
| Max. Fill Time | 100ms | |
| Lock Mass | m/z 445.120025 | Dodecamethylcyclohexasiloxane [3] |
| Scan Range | m/z 395-1500 | |
| Picotip Needle | 20µm / 10µm | NewObjectives, Ithaca, USA |
| Voltage | 2.3-2.7kV | (might vary between experiments) |
| **MS2 High Res** | **Top10** | **HCD** |
| Resolution | R15000 at m/z 200 | |
| AGC | 1E5 | |
| Max. Fill Time | 50ms | |
| Isolation | 2.0 m/z | |
| Isolation window Offset | 0.3 m/z | |
| Scan Range | 200-2000 m/z | |
| Fixed 1st Mass | - | |
| Norm. Collision Energy | 27 | |
| Threshold Int. / Min AGC Target | 2E3 / 4E4 | |
| Charge states | Unassigned, 1, 6-8, >8 | (rejected) |
| Dynamic Exclusion | 15s / 3ppm | |

TABLE S1. Mass Spectrometry instrumentation parameters for Q-exactive HF

| Instrument / Material | Manufacturer (Supplier) | Comments |
|---|---|---|
| **Dionex3000 RSLC** | ThermoScientific, Idstein, Germany | Nanoflow System |
| Acclaim PepMap 100 C18, | | Trap-Column Setup |
| 3 µm, 300 µm x 5 mm, | ThermoScientific, Idstein, Germany | Load: 2µl/min |
| Acclaim PepMap C18 2 µm, 75 µm x 15 cm | | Separation: 200nl/min |
| Picotip Needle 20 µm / 10 µm | NewObjectives, Woburn, USA | |

TABLE S2. Mass Spectrometry instrumentation parameters for Thermo Dionex3000 RSLC

| Instrument / Material | Manufacturer (Supplier) | Comments |
|---|---|---|
| MASCOT V2.6 [4] | MatrixScience, London, UK | Protein Identification Software<br>matrixscience.com |
| MSConvert V3.0 [2,5] | Proteowizard, CDN | File Conversion Tool |
| Progenesis QIP V4.2 | Nonlinear Dynamics, Newcastle u.T., UK | Quantitative data interpretation (Peak Picking, MS/MS Export,<br>Peptide ID import and Assignment, Quantification (MI3/Hi3)) |
| Scaffold V4.11 [6] | Proteome Software, Portland, OR, USA | Protein Identification Statistics and Validation, Visualization<br>Proteomesoftware.com |

TABLE S3. Mass Spectrometry software versions used for analysis

| **Mascot Parameter QE-HF** | Value | Comments |
|---|---|---|
| **Version** | 2.6 | |
| MS Tolerance | 10 ppm | |
| Protease | Trypsin | |
| Missed Cleavages | 3 | |
| Fixed Modifications | | |
| Variable Modifications | Acetyl- N-Protein, Oxidation (M) | |
| MS/MS Tolerance | 30 mmu | |
| Instrument | ESI-Quad | |
| Databases | Contaminants<br>Enzymes<br>STDs_TAGs | Databases applied according to sample origin. |
| Decoy | Yes | |

TABLE S4. Mass Spectrometry Mascot software parameters

| **Item (Chemicals)** | Manufacturer (Supplier) | Order Number |
|---|---|---|
| NH4HCO3 Ammoniumbicarbonate | Sigma | A-6141 |
| Water HPLC grade | Merck, Darmstadt, Germany | 1.15333.2500 |
| Acetonitrile HPLC Grade | Merck, Darmstadt, Germany | 1.00029.2500<br>1.00030.2500 |
| Formic Acid p.a. | Merck, Darmstadt, Germany | 1.00264.0100 |
| Trypsin Gold sequencing grade, (modified Trypsin) | Promega, Walldorf, Germany | V5280 |

TABLE S5. Mass Spectrometry chemicals used
